# Genomic analysis of cultivated infant microbiomes identifies *Bifidobacterium* 2’-fucosyllactose utilization can be facilitated by co-existing species

**DOI:** 10.1101/2023.03.10.532136

**Authors:** Yue Clare Lou, Benjamin E. Rubin, Marie C. Schoelmerich, Kaden DiMarco, Adair L. Borges, Rachel Rovinsky, Leo Song, Jennifer A. Doudna, Jillian F. Banfield

## Abstract

Human milk oligosaccharides (HMOs) ensure proper infant gut microbiome establishment. Isolate studies have revealed the genetic basis for HMO metabolism, but they exclude the possibility of HMO assimilation via synergistic interactions involving multiple organisms. Here, we investigated microbiome responses to 2’-fucosyllactose (2’FL), a prevalent HMO and infant formula additive, by establishing individualized microbiomes using fecal samples from three different infants as the inocula. *Bifidobacterium breve*, a prominent member of infant microbiomes, typically cannot metabolize 2’FL. Using metagenomic data, we predicted that extracellular fucosidases encoded by co-existing members such as *Ruminococcus gnavus* initiate 2’FL breakdown, thus critical for *B. breve’s* growth. Using both targeted co-cultures and by supplementation of *R. gnavus* into one microbiome, we show that *R. gnavus* can promote extensive growth of *B. breve* through the release of lactose from 2’FL. Overall, microbiome cultivation combined with genome-resolved metagenomics demonstrated that HMO utilization can vary with an individual’s microbiome.

## Introduction

The early-life gut microbiome is a simple yet rapidly changing ecosystem crucial for infant development^1–3^. Early-life events, such as feeding, influence the succession of the gut microbiome^2,4,5^. Breast milk is considered the preferred food source for infants due to its protection against infections and allergy development, among other benefits^6–11^. Many of these advantages can be attributed to human milk oligosaccharides (HMOs), a group of complex carbohydrates unique to human milk that can only be metabolized by certain gut commensals^12,13^. While breast milk is ideal, it is not accessible to all infants. Within the first 6 months of life, over 70% of infants are estimated to receive formula^14,15^, which lacks key breast milk compounds, including HMOs, and can potentially lead to negative health outcomes^16,17^.

HMO supplementation is becoming a common practice for shortening the functional gap between infant formula and breast milk^16^. Of the over 200 different HMOs that have been identified in breast milk, a few have been industrially produced and added to infant formula^18^. One of these is the α-1,2-fucosylated trisaccharide 2’-fucosyllactose (2’FL). 2’FL is one of the most abundant and prevalent HMOs in breast milk^19,20^. Its presence in feeding has been positively associated with infant health^21^. However, such benefits can be infant-specific, as variations in 2’FL utilization exist between infant gut microbiomes^22–24^. Compositional differences in *Bifidobacterium* spp. partly explain microbiome-specific responses to 2’FL^25^ as different *Bifidobacterium* species have varying abilities to metabolize HMOs^26^. Further, variations among strains of the same *Bifidobacterium* species also contribute to interpersonal differences in 2’FL utilization. For instance, only a minority of strains of *Bifidobacterium breve*, an abundant infant gut colonizer, can break down 2’FL^26–29^. Given the natural prevalence and commercial application of 2’FL, we sought to elucidate additional mechanisms that explain microbiome-specific responses to 2’FL.

Prior studies examining 2’FL metabolism mainly focused on *Bifidobacterium*, the primary HMO degrader^26^. 2’FL utilization by co-existing *Bifidobacterium* species suggests the importance of microbial interactions in HMO metabolism^26,30–32^. Besides *Bifidobacterium*, some other community species may also encode genes for 2’FL breakdown, and thus could influence *Bifidobacterium’s* growth on 2’FL^33,34^. In part due to the lack of experimentally tractable gut microbiomes, it remains largely underexplored how co-existing members interact with *Bifidobacterium* in 2’FL assimilation. To investigate this, we established relatively representative gut microbiomes from biologically unrelated preterm and full-term infants. As we created consortia comprised of natural populations recovered directly from human infant stool samples, our work differs from prior *in vitro* microbiome research and enabled person-specific microbiome analyses^35,36^. By leveraging functional predictions from metagenomics analyses and testing these predictions experimentally using cultivated microbiomes, we showed that metabolic functions performed by certain co-existing members could influence the growth of *B. breve*,which cannot utilize 2’FL on its own, in 2’FL-supplemented media. Our work expands the current understanding of individualized microbiome responses to 2’FL and provides a testing platform for infant-specific nutrient responses, which could, in turn, be leveraged for more targeted and effective infant feeding regimes.

## Results

### Infant gut microbiome cultivation scheme

To elucidate how community interactions may influence *Bifidobacterium*’s growth on HMOs, we selected infant gut microbiomes as inocula based on the presence of a single *Bifidobacterium* species^37^. Microbiomes from three biologically unrelated, primarily breastfed infants were chosen, each of which had a distinct composition and complexity, yet they shared a near-identical *Bifidobacterium breve* strain (~99.6% genome-wide average nucleotide identity (gANI)) (Methods). The lack of multiple *Bifidobacterium* species enabled the examination of interactions between *non-Bifidobacterium* and *Bifidobacterium* involved in HMO metabolism.

To explore growth conditions that would allow the maximal recovery of the inoculum species, we began our cultivation efforts with one stool sample, collected from a three-month-old full-term infant, which we named “FT-1.” Its microbiome consisted of 8 bacterial species across three phyla (Actinobacteria, Firmicutes, and Proteobacteria) that were ≥0.1% abundant (Figure S1).

To identify a growth medium that allowed the recovery of a microbiome compositionally as close to the original community as possible, we inoculated FT-1 in five complex media (Figure 2A). Brain-Heart Infusion (BHI) was selected as the primary base since this medium allowed the isolation and stable maintenance of adult gut microbiomes in previous studies^36,38^. We also included the modified Gifu anaerobic medium (mGAM), another rich medium used to isolate individual gut microbes and consortia^39,40^. Given that the FT-1 infant was fed with human breast milk (HM) primarily, we also included HM as a growth medium. Mucin was supplemented since it can be a key carbon source for some gut species such as *Ruminococcus gnavus* in FT-1^41^. Cultures were inoculated with phosphate-buffered saline (PBS)-resuspended liquid stool and anaerobically passaged in a 48-hour interval for three passages. Enrichments were harvested after the 1st and the 3rd passaging, and their compositions were assessed via metagenomics sequencing (Methods) (Figure 1).

**Figure 1.**
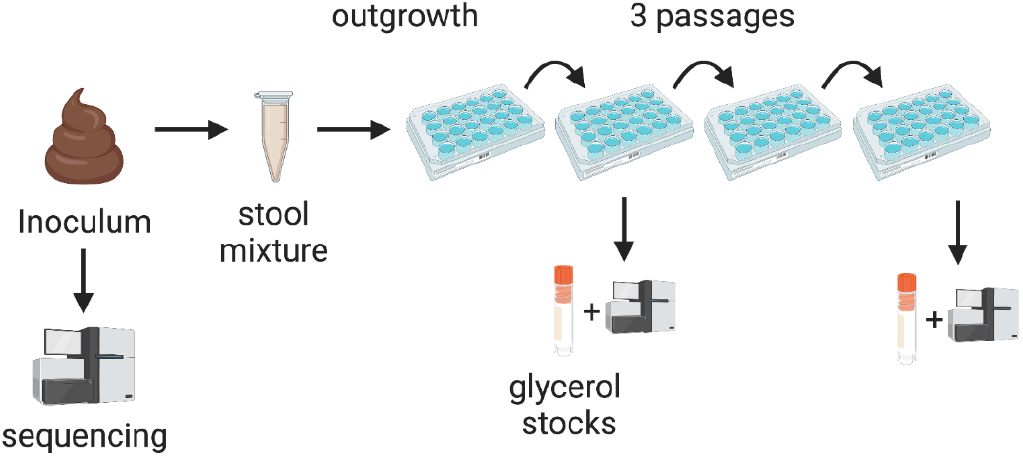
Microbiome cultivation experimental setup. The frozen pilot infant fecal sample was resuspended in a PBS solution before inoculating it into anaerobic media. After an initial 48-hour outgrowth period, the enrichments were subsequently passaged every 48 hours for a total of three passages. A fraction of the enrichment was harvested from the 1st and the 3rd passaged enrichments for sequencing and long-term storage.

**Figure 2.**
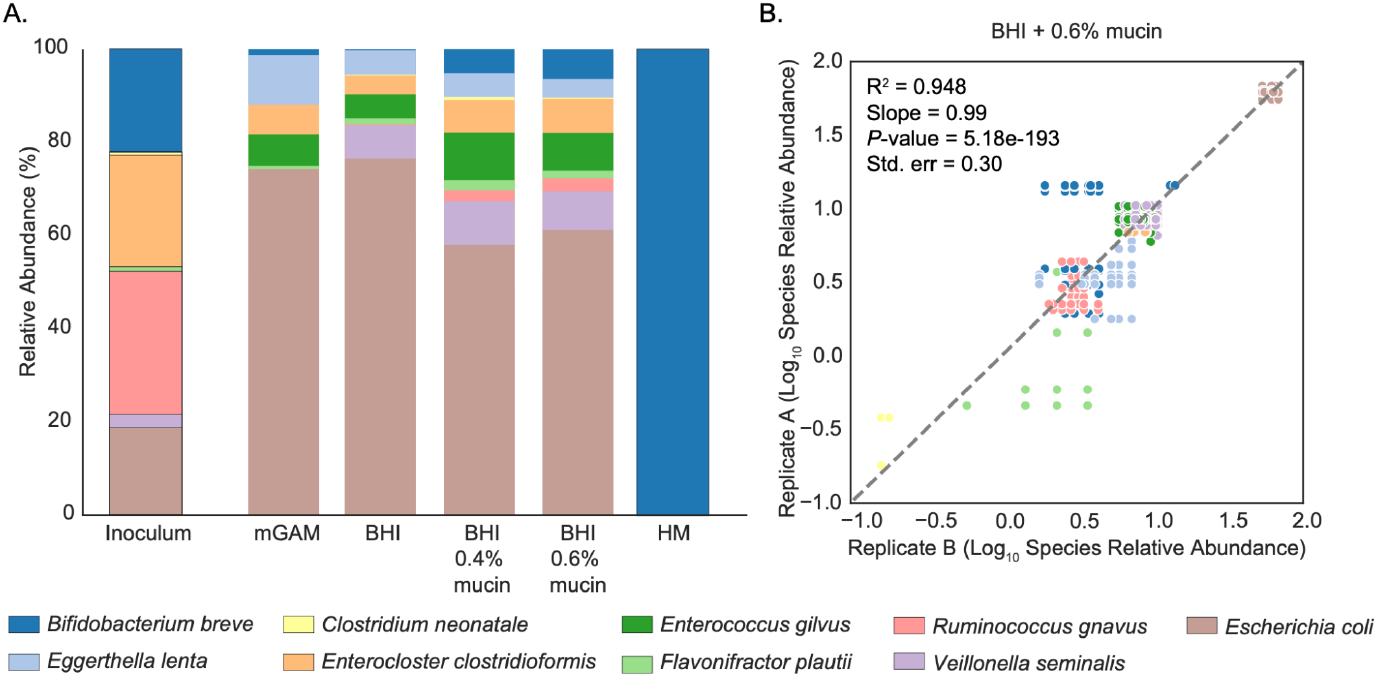
FT-1 microbiome cultivation overview. A) Bar height represents normalized species relative abundance, and bars are colored by species. The x-axis represents the growth media, and the data are the averages of replicates (n= 2 or 3). B) Pairwise species relative abundance comparison among different replicates. Correlations were calculated using the two-sided linear least-squares regression.

### BHI with mucin addition recovered a representative and stable infant gut microbiome

For all growth media, a compositionally stable community was largely reached after just one passage (Figure S2). Except for HM, a near-complete microbiome was recovered (Figures 2A and S2). HM, intriguingly, resulted in the least diverse community in which *Bifidobacterium breve* was the only species recovered (Figure 2A). The lack of detectable organisms in HM-only negative controls indicated *B. breve* was not derived from HM. Additional passaging of the *B. breve*-only HM enrichments in BHI + 0.6% mucin (wt/vol) did not change the community composition (Figure S2B), suggesting *B. breve* was the only viable species in HM enrichments.

Of the non-HM conditions, mGAM had the lowest microbiome recovery rate (Figure 2A). All species were recovered in at least one replicate of BHI, BHI + 0.4% mucin, or BHI + 0.6% mucin, yet, the mucin supplementation yielded a community more structurally similar to the inoculum than the BHI-only medium (p = 0.034; Wilcoxon rank-sum test on weighted UniFrac Distances). Of the species in the inoculum, *Clostridium neonatale*, present in about a third of the replicates of the BHI-based media with an average abundance of ~1%, had the lowest replicability (Figure 2A). The stochastic presence of *C. neonatale* in enrichments could be partly due to its low abundance in the inoculum (0.54%; Figure 2A). In contrast, *E. gilvus*, undetectable in the initial inoculum, was detected in nearly all non-HM conditions (Figure 2A). Given its prevalence in enrichments and absence in reagent-only negative controls, we speculated that this species was indeed present in the original stool microbiome but was too low in abundance to be detected. This led us to conclude the FT-1 microbiome was, in fact, a 9-species community.

Communities grown in non-HM media were dominated by *Escherichia coli*. This was particularly evident in media with no mucin supplementation, in which *E. coli* abundance increased 4-fold compared to the inoculum, reaching ~75% (Figure 2A). We hypothesized that adding polysaccharides, such as mucin, could give a growth advantage to other community members such as Firmicutes since they can metabolize complex carbohydrates whereas *E. coli* cannot^42,43^. Indeed, mucin supplementation significantly lowered *E. coli* abundance (q = 0.0021; Wilcoxon rank-sum test) (Figure 2A). The drop in *E. coli* was accompanied by an increased abundance of bacteria from Actinobacteria and Firmicutes (Table S1). While both 0.4% and 0.6% mucin concentrations yielded compositionally stable and reproducible enrichments, 0.6% mucin supplementation resulted in a more stable growth of *B. breve* than 0.4% mucin (Figures 2B and S2; Table S2) (Methods). BHI + 0.6% mucin was thus selected as the base medium for subsequent experiments involving FT-1.

### Near-complete and stable *in vitro* gut microbiomes from two additional infants

To test the more general value of BHI-mucin media for community cultivation, we inoculated the other two infant fecal samples into BHI with either 0.4% or 0.6% mucin (Methods). One sample was collected from a 56-day-old full-term infant, which we named “FT-2”, and the other sample was from a 15-day-old preterm infant, which we named “PT-1”. Both infants were mostly breastfed. The FT-2 microbiome consists of 20 bacterial species from Actinobacteria, Firmicutes, and Proteobacteria phyla. The PT-1 community, consisting of six bacterial species, was dominated by Proteobacteria, a typical preterm gut microbiome structure^37,44^ (Figure S1).

Like FT-1, some organisms that were low in abundance in FT-2 and PT-1 fecal samples were enriched in the cultivated communities (Figures 3 and S3). In addition, in BHI-mucin media inoculated with FT-1, FT-2, or PT-1, the abundance of Actinobacteria decreased relative to that in the fecal sample (Figures 2A, 3, S2, and S3). Each of the cultivated microbiomes only had one Actinobacteria strain, which was *B. breve*. While *B. breve* represented ~3% and ~5% of FT-1 and FT-2 BHI-mucin enrichments, respectively, this species was nearly undetectable in PT-1 enrichments. Intriguingly, given their different initial compositions, inoculating these three microbiomes into HM led to enrichments containing only *B. breve* (Figures 2A, 3, S2, and S3).

**Figure 3.**
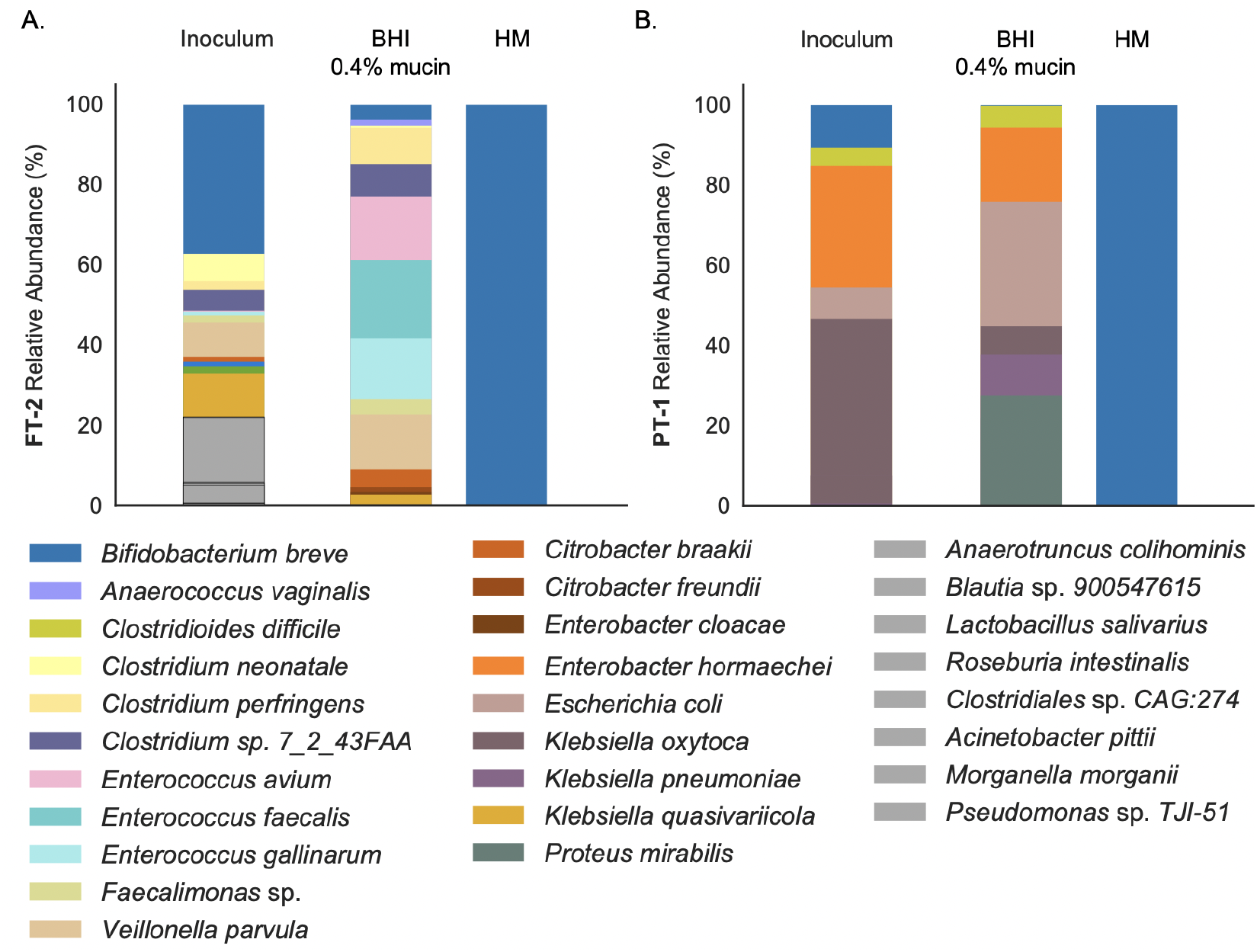
Inoculation of three additional infant gut microbiomes. Average compositions of the FT-2 (A) and PT-1 (B) enrichments grown in BHI+0.4% mucin and HM. Bar height represents normalized species relative abundance, and bars are colored by species. Organisms colored in gray (N=8) were present in the FT-2 stool inoculum but not in any cultivated communities. All organisms in PT-1 were detected in cultivated communities. The x-axis represents the growth media.

Overall, 60% and 100% of the FT-2 and PT-1 inocula species were recovered in BHI + 0.4% mucin, respectively, higher than the microbiome recovery rates of BHI + 0.6% mucin. Further, replicability was higher when the BHI + 0.4% mucin media was used compared to the BHI + 0.6% mucin media (Figure S3A and B). Given that the FT-2 and PT-1 community compositions remained largely unchanged in BHI + 0.4% mucin over three passages (Figure S3C), this medium was selected for subsequent experiments for these two infant microbiomes.

### HM exerts strong selective pressure on the infant gut microbiome

The difference in the abundance of *B. breve* in HM and BHI-based media motivated us to test five additional media that contained HM and BHI mixed in different volume proportions (Figures 4 and S4). Since all three infant gut microbiomes had similar growth outcomes in HM versus non-HM media, FT-1 was selected as the representative for investigating HM’s effects on the gut consortia. Cultures were passaged once before community composition profiling (Methods).

**Figure 4.**
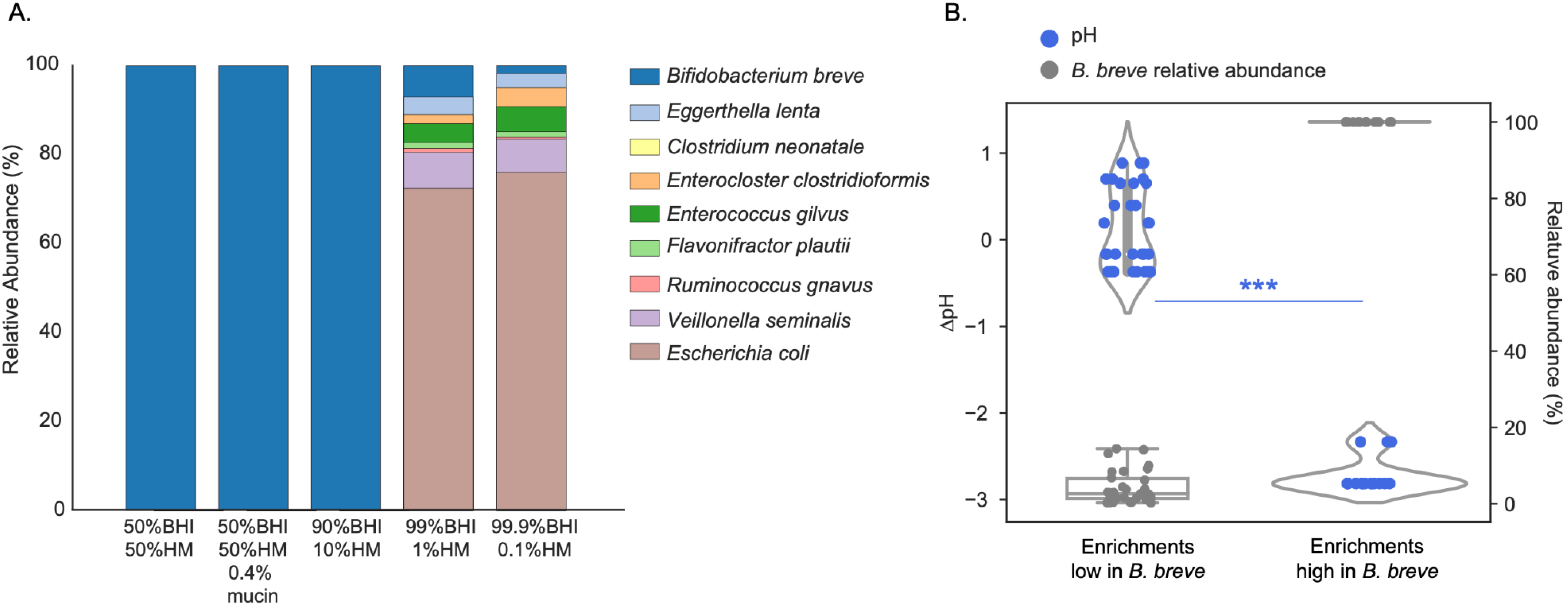
HM poses strong selective pressure on the FT-1 microbiome. A) Bar height represents normalized species relative abundance, and bars are colored by species. The x-axis represents the growth media, and the data are averages of the replicates. B) The left y-axis represents the change in pH, and the right y-axis represents the relative abundance of *B. breve*. Each circle represents a culture replicate and is colored by either ΔpH (blue) or *B. breve* abundance in that culture (gray). The x-axis represents the community type, either dominated or not dominated by *B. breve* (*** *p* < 0.001).

We found that HM, even a small quantity, could strongly affect the FT-1 microbiome composition. With as little as 10% (vol/vol) HM in growth media, *B. breve* remained to be the only species of the enrichments. When the HM concentration dropped below 1%, *B. breve* abundance decreased, and the enrichment community diversity increased (Figure 4A). Interestingly, enrichments grown in BHI with ≤1% HM (99% BHI+1% HM and 99.9% BHI + 0.1% HM) were compositionally similar to those grown in BHI with mucin supplementation (BHI + 0.4% mucin and BHI + 0.6% mucin) (Figures 2A and 4A).

HM’s influence on FT-1 was mostly driven by *B. breve* abundance. As HM concentrations increased, *B. breve* was significantly enriched (Figures 2, 4, and S4) (Spearman correlation = 0.86, *p* = 1.37e-11). Notably, communities became increasingly acidic with rising *B. breve* abundance (Spearman correlation = −0.88, *p* = 1.29e-12). The changes of pH in *B. breve*-dominated communities (*B. breve* relative abundance >50%) were significantly larger than in communities in which *B. breve* was at low abundance (*p* = 1.92e-11; Wilcoxon rank sum test) (Figure 4B).

### 2’FL exerts infant-specific effects on microbiome compositions

Having shown that HM could profoundly shape the infant gut microbiome by promoting the growth of *B. breve*, an effect likely mediated through milk carbohydrates including HMOs, we next investigated if a single HMO, 2’FL, would recapitulate this effect. We included a 2’FL concentration roughly equivalent to that in human milk (0.3% (wt/vol))^13^ and a 10x lower one (0.03%) to see if there was a strong concentration-dependent effect on the microbiomes. Communities were grown in BHI-mucin-based media with one passage, as described above (Methods).

When 0.03% 2’FL was added, all enrichments resembled those grown in BHI-mucin-only media (Figures 2, 3, and 5A-C). When 0.3% 2’FL was supplemented, only the FT-1 community composition changed dramatically (Figure 5). Comparing 0.3% to the 0.03% 2’FL supplementation, *B. breve* abundance in FT-1, which had a 33% decrease in species richness, increased from ~6% to ~36% (Figures 5A, D, and S5; Table S3). Such a community structure remained mostly unchanged when growing the FT-1 community in media supplemented with 2’FL concentrations higher than 0.3% (Figures 5F and S5). Notably, when removing glucose, whose concentration is 0.3% (wt/vol) in the BHI used for community cultivation, the abundance of *B. breve* significantly increased (q = 0.025; two-sided Welch’s t-test), reaching ~72%, and *E. gilvus, E. coli*, and *V. seminalis* significantly decreased in abundance (q < 0.05; two-sided Welch’s t-test) (Figure S6; Table S4). The community diversity of FT-2 remained the same regardless of 2’FL concentrations, although the abundances of several species, including *B. breve*, changed significantly (Figure 5B and E; Table S3). The community composition and diversity of PT-1 stayed the same across different 2’FL concentrations (Figure 5C).

**Figure 5.**
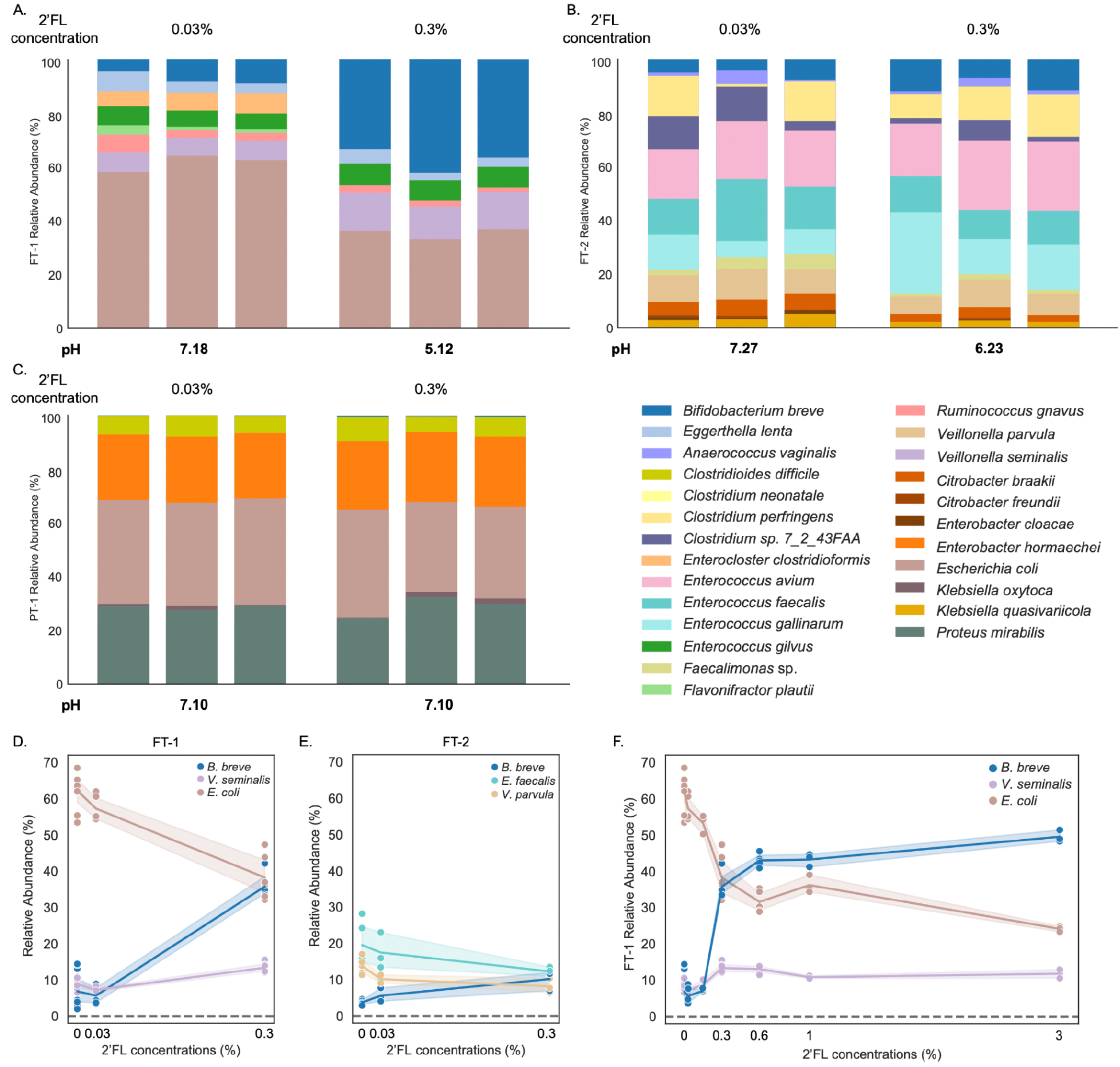
2’FL supplementation resulted in distinct responses from different infant gut microbiomes. A-C) The compositions of the FT-1 (A), FT-2 (B), and PT-1 (C) enrichments grown in BHI-mucin supplemented with 0.03% and 0.3% 2’FL. Bar height represents normalized species relative abundance, and bars are colored by species. The x-axis represents the growth media, and all replicates are shown. The pH of each enrichment is the mean across the three replicates. The figure legend is shown on the bottom right of the figure panel. D-E) The significant enrichment or depletion of FT-1 (D) and FT-2 (E) species cultured in different 2’FL concentrations (0, 0.03%, and 0.3%). Each circle represents a species whose abundance changed significantly in different 2’FL concentrations (Spearman correlation |r| ≥ 0.8 and q < 0.05; Table S3). The coloring scheme is the same as the one used in the stacked bar plots. The x-axis represents the 2’FL concentrations. F) The FT-1 inoculum grown in media with up to 3% 2’FL and species that showed a significant depletion or enrichment in Figure 5D are shown. The coloring scheme is the same as the one used in the stacked bar plots. The x-axis represents 2’FL concentrations.

The changes in community compositions in response to 2’FL concentrations were also reflected in the pH. In media that contained 0.03% 2’FL, the pH values of all enrichments were ~7.1. However, with a 2’FL concentration of 0.3%, the pH of the FT-1 and FT-2 enrichments dropped by 2.1 and 1.0, creating an acidic milieu, whereas the pH of PT-1 remained the same (Figure 5A-C). Based on these findings, we infer that the effects of 2’FL at the tested concentration levels can be infant-specific. In some cases, it can lead to a significant enrichment of gut commensals such as *B. breve*.

### In FT-1, *B. breve* could utilize 2’FL in the presence of *R. gnavus*

To elucidate why 2’FL supplementation resulted in distinct effects on these infant microbiomes, we assessed the genomically defined metabolic potentials of the communities, focusing primarily on FT-1 because it exhibited the most substantial change at the physiologically relevant 2’FL concentration of 0.3%. All genomes in FT-1 were reconstructed, and functional predictions were made using CAZyme, KEGG, Pfam, TCDB, and signalP (Methods).

We first examined whether *B. breve* in FT-1 could directly metabolize 2’FL by isolating it from FT-1 and growing it in media with 2’FL as the sole carbohydrate source (Methods). No growth was detected (Figure S7). Genomic analysis revealed that *B. breve* encoded one fucosidase, GH95, but this gene was predicted to be intracellular due to the lack of a signal peptide. With the lack of 2’FL transporters, we concluded that *B. breve* from FT-1 cannot metabolize 2’FL (Methods).

Since *B. breve* cannot break down 2’FL, we hypothesized that the *B. breve*-promoting effect of 2’FL seen in FT-1 resulted from other community members liberating lactose and fucose from 2’FL, which could subsequently be metabolized by *B. breve*. A community-wide CAZyme survey revealed that *R. gnavus* was the only organism besides *B. breve* in FT-1 whose genome encodes fucosidases (Methods). Specifically, it encodes six fucosidase genes (three GH29 and three GH95 genes), two of which (one GH29 and one GH95 gene) were predicted to be extracellular. To investigate whether the presence of *R. gnavus* was sufficient for *B. breve* to grow on 2’FL, we isolated *R. gnavus* from FT-1 and inoculated this strain as well as the *B. breve* strain isolated from FT-1 as mono- and co-cultures in media with 2’FL or its constituents, L-fucose or D-lactose, as the sole carbohydrate source (Methods). The growth was measured using optical density (OD, λ = 600 nm), and the coculture compositions were assessed via amplicon sequencing (Methods).

The *R. gnavus* monoculture grew on 2’FL, but not the *B. breve* culture (Figure 6A). The co-culture had a higher growth rate and yielded a higher final cell density than the *R. gnavus* monoculture (Figure 6A; Table S5). Notably, *B. breve* dominated the co-culture community (~80% of the community) (Figure 6A right panel), paralleling the findings of the FT-1 community grown on 2’FL (Figure 5A).

**Figure 6.**
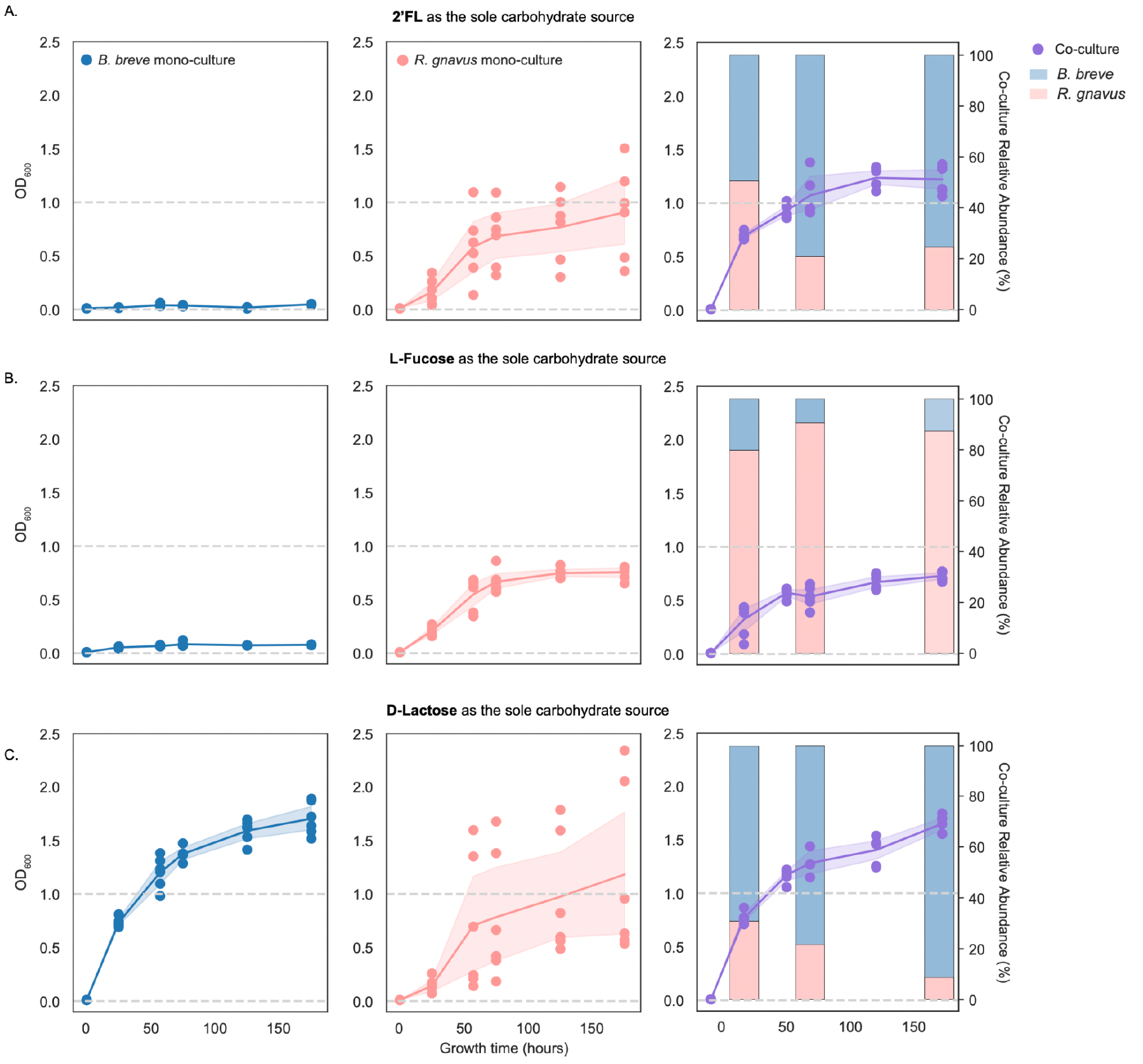
Mono- and co-culture growth on 2’FL, L-fucose, and D-lactose. *B. breve* (blue) and *R. gnavus* (pink) monocultures and cocultures (purple) grew in media in which 2’FL (A), L-fucose (B), or D-lactose (C) was the only carbohydrate source. The x-axis represents the growth time (in hours). The left y-axis represents OD_600_, and the right y-axis in the co-culture panels represents the normalized relative abundance, calculated using sequencing coverage. Each circle represents a mono- or co-culture sample. The relative abundance of each species (*B. breve* or *R. gnavus)* at the 24^th^, 72^nd^, and 168^th^ hours was shown in stacked bar plots in less saturated colors in the co-culture panels.

*R. gnavus* but not *B. breve* could grow on L-fucose (Figure 6B). The growth rate and the final cell density of the co-culture were similar to those of the *R. gnavus* monoculture grown on fucose (Figure 6B; Table S5). In contrast to growth on 2’FL, this co-culture was dominated by *R. gnavus* (~80% of the community) (Figure 6B right panel).

Both the *R. gnavus* and *B. breve* monocultures grew on D-lactose (Figure 6C). *B. breve* had a higher growth rate and reached a higher final cell density than *R. gnavus* (Figure 6C; Table S5). The co-culture’s growth rate and the final cell density were comparable to those seen in the *B. breve* monoculture (Figure 6C; Table S5), and the community was dominated by *B. breve* (Figure 6C right panel).

Overall, the experiments revealed that *R. gnavus* grows on 2’FL, lactose, and fucose, whereas *B. breve* can only grow on lactose. Given that *B. breve* did not show extensive growth on 2’FL or fucose, we infer that *B. breve’s* growth in the co-culture grown on 2’FL was due to the lactose released by *R. gnavus*. As the only member that can break down 2’FL, *R. gnavus* is likely an essential FT-1 member when grown on 2’FL. The community grown on ≥0.3% 2’FL was dominated by *B. breve* and *E. coli*, followed by *V. seminalis, E. gilvus, E. lenta*, and last, *R. gnavus* (Figure 5A). We thus infer that *R. gnavus* is a keystone member. *E. coli* has the potential to metabolize lactose and fucose, *E. gilvus* is predicted to metabolize lactose, and *V. seminalis* likely metabolizes fucose. *E. lenta* did not encode genes for metabolizing either sugar, suggesting it potentially utilizes by-products from the FT-1 community (Methods) (Table S6).

### Supplementation of *R. gnavus* and 2’FL shifts a preterm microbiome into a full-term-like, *Bifidobacterium-enriched* community

Given the dependency of *B. breve* on *R. gnavus* in FT-1 grown on 2’FL, we sought explanations for the extensive growth of *B. breve* in FT-2 and the limited growth in PT-1. The presence of the intracellular GH95 fucosidase and the lack of 2’FL transporters suggest none of the *B. breve* strains in these inocula could directly utilize 2’FL. Thus, we applied the same genome-resolved method used for FT-1 to the FT-2 and PT-1 communities (Methods).

Three species *(Clostridium perfringens, Clostridium* sp. 7_2_43FAA, and *Faecalimonas* sp.) in FT-2 are predicted to metabolize 2’FL extracellularly (Methods) (Table S7). Since the growth of *B. breve* on 2’FL was less significant in FT-2 than in FT-1, we speculated that other factors, such as inter-species competition, might result in limited *B. breve* growth. The extensive growth of *B. breve* in FT-1-based mono- and co-cultures, when grown in media with lactose as the sole carbohydrate (Figures 6A and C), led us to hypothesize that other organisms in FT-2 might be directly competing against *B. breve* for lactose. Indeed, we found that the genomes of the three extracellular fucosidase-encoding species also encode extracellular β-galactosidases (Table S7), some of which are genomically adjacent to the extracellular fucosidases, suggesting these organisms could further break down extracellular lactose freed from 2’FL into glucose and galactose. No extracellular β-galactosidases were detected in the FT-1 or the PT-1 community. We thus infer that the lower abundance of *B. breve* in FT-2 compared to FT-1 in 2’FL-supplemented media may be the consequence of the degradation of lactose, *Bifidobacterium’s* preferred carbohydrate source^45,46^, by other community members, which reduces the amount of lactose that can be used by *B. breve* for growth.

In PT-1, no other organisms besides *B. breve* encode a fucosidase. This explains why the PT-1 community failed to respond to 2’FL supplementation and remained to be dominated by Proteobacteria (Figure 5C). To test whether adding a strain encoding extracellular fucosidases would be sufficient to enrich *B. breve* in PT-1 when sufficient 2’FL is supplemented, we inoculated *R. gnavus* isolated from FT-1 to PT-1 (Methods). Notably, *B. breve* was significantly enriched in the *R. gnavus*-positive PT-1 microbiome grown in media with 0.3% 2’FL and reached an abundance level similar to that seen in FT-1 (Figures 5A and 7). As in the FT-1 enrichment, the increase in *B. breve* abundance in the *R. gnavus*-supplemented PT-1 microbiome was correlated with a decrease in community species diversity and pH, reaching an average value of 5.15, ~1.8 points lower than the unsupplemented PT-1 microbiome grown under the same conditions (Figure 7).

**Figure 7.**
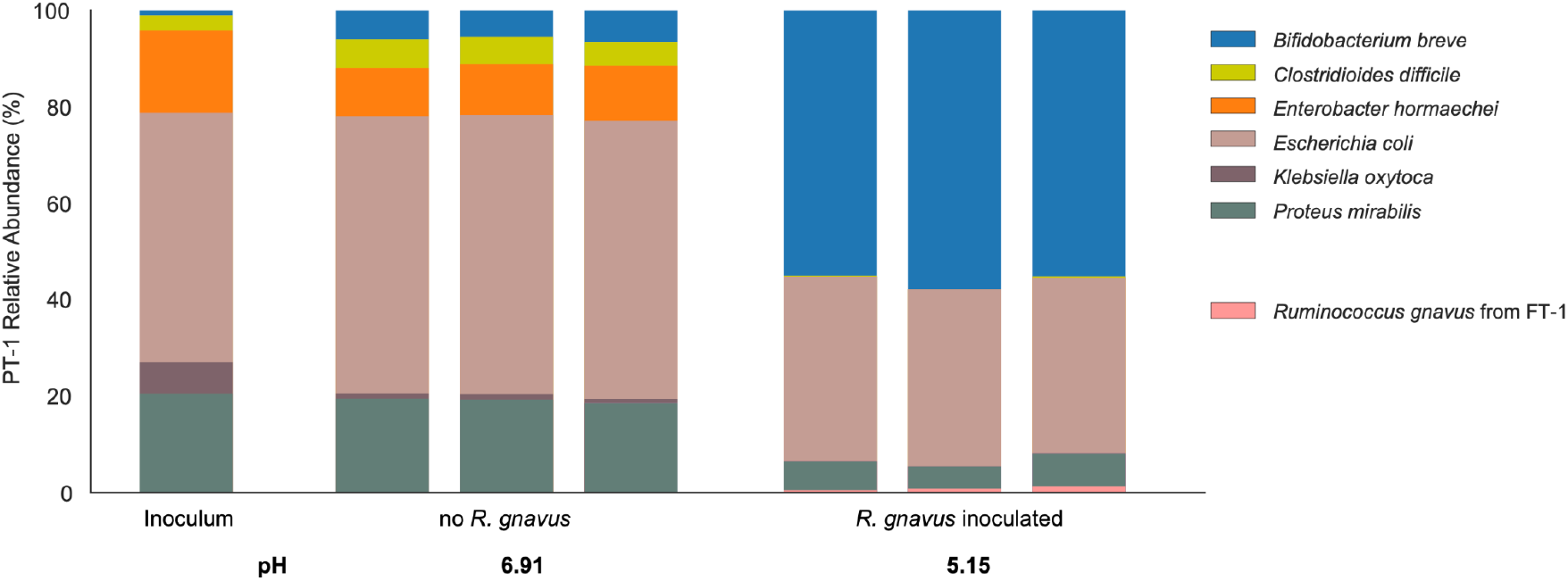
The addition of *R. gnavus* to PT-1 enriched *B. breve* in 2’FL-supplemented media.

The PT-1 community with or without the supplementation of *R. gnavus* was grown in media containing 0.3% 2’FL. Bar height represents normalized species relative abundance, and bars are colored by species. The x-axis represents the community type, and all replicates are shown.

## Discussion

Human health is directly impacted by the gut microbiome^47^, whose composition and function are shaped by microbial interactions^48–50^. Despite the importance of gut consortium interactions, their consequences are understudied. This is in part due to the lack of tractable gut microbiome models^51^. Experimental microbiome perturbations are usually impossible in humans, and large-scale observational studies are confounded by host-associated factors, such as host genetics and lifestyle, making it difficult to identify causative microbial mechanisms for a given phenotype^52^. Animal models are tractable compared to humans but are expensive, labor-intensive, and have relatively low throughput^53^. Complex continuous culture systems mimicking the human gastrointestinal system have revealed key insights into the gut microbiome ecology but have limited throughput as well^54,55^. It is therefore important to establish reasonably representative human microbiome model systems that can be used to explore microbial interactions and their consequences over a wide range of conditions. The infant gut microbiome, compositionally simple yet distinctive for each person^37^, represents a unique system for experimentally investigating gut microbiome interactions and their functional consequences. Here, using clinically relevant stool samples of human infants, we established stable, reproducible, and person-specific laboratory gut consortia. We showed that supplementing complex carbohydrate sources such as mucin or human milk (HM) could limit the bloom of Enterobacteriaceae, a common community cultivation issue^36,40^, by promoting the growth of other taxonomic groups, resulting in *in vitro* consortia with fairly representative structures. Since these supplemented components are common in the infant gut, our work demonstrates the importance of using microbiome cultivation media that closely mimic the nutrient content of the actual environment.

Diet affects gut microbiome compositions and functions^1,56^. As each human is colonized by a distinct gut microbiome, it is important to study dietary effects on different microbiomes to uncover the diversity of relevant interactions and commonalities across individuals^57,58^. To identify infant-specific microbiome responses to common perturbations, we established *in vitro* consortia using stool samples as inocula. Our top-down approach differs from, but complements, prior bottom-up microbiome research using generic synthetic communities made of prevalent isolates^35^. Preserving person-specific microbiome signatures, the direct-stool-resuspension approach has been previously applied by others to examine personalized adult gut microbiome drug metabolism^40,59^. In this study, we investigated microbiome-specific utilization of common infant nutrient sources. By comparing cultivation outcomes and functional potentials of different gut microbiomes, we identified crucial microbial functions, which can be carried out by taxonomically distinct organisms in different infants, for metabolizing human milk carbohydrates.

We found that *B. breve*, regardless of the initial abundance and community composition, reached nearly 100% abundance in HM enrichments. To our knowledge, this is the first study in which human milk is directly used to cultivate infant gut microbiomes. By experimentally showing a *B. breve*-dominant community can be established using media containing as little as 10% HM, we demonstrated the significant role HM plays in shaping gut microbiome compositions. The enrichment of *B. breve* in HM-based media aligns with findings from observational studies that breastfeeding results in a *Bifidobacterium-enriched* gut microbiome with a lower species diversity when compared to that of formula-fed infants^4,60,61^. The bifidogenic effect of HM is partly due to *Bifidobacterium* efficiently fermenting breast milk carbohydrates, creating a low-pH environment unsuitable for the growth of many members including potential pathogens (i.e., Proteobacteria)^62,63^. Indeed, consistent with prior *in vivo* studies^27,32,64–68^, in our enrichments, we also observed a negative correlation between *B. breve* abundance and the pH, as well as Proteobacteria abundance. The near-complete dominance of *B. breve* in HM enrichments slightly deviates from the natural gut microbiomes of breastfed infants. We speculate the greater prevalence of *B. breve* in HM enrichments compared to the infant gut is due to the higher concentration of lactose, which is mostly absorbed in the infant’s small intestine^69^. Using *B. breve* isolated from an infant gut consortium, we showed that this species could efficiently use lactose and grow on it as the sole carbohydrate source. Further supporting our hypothesis is the observation that extracellular β-galactosidases-encoding species, which degrade lactose, could limit the growth of *B. breve*. Overall, our cultivated microbiomes largely recapitulated the bifidogenic *in vivo* responses to breastfeeding. What makes this microbiome system powerful is the ability to directly manipulate gut communities, allowing the identification of potential phenotype-causing microbial mechanisms.

The significant enrichment of *B. breve* in HM indicates that this species can efficiently utilize HM compounds for growth. Besides lactose, HMOs are the most abundant carbohydrates in HM^12,13^. Unlike lactose, HMOs are mostly unabsorbed by infants and are instead exclusively metabolized by some gut commensals including *Bifidobacterium^12,13^*. In this work, we examined the growth of *B. breve* and the rest of the community in media supplemented with 2’-fucosyllactose (2’FL), one of the most abundant HMOs^70^. All microbiomes studied here share a near-identical *B. breve* strain that cannot break down 2’FL. However, we showed that it could grow well so long as it co-exists with species that encode extracellular fucosidases. This is particularly evident with the Proteobacteria-dominant preterm gut microbiome (PT-1), which lacks community members encoding extracellular fucosidases and showed minimal response to 2’FL, including no detectable growth of *B. breve*. However, when supplemented with extracellular fucosidase-encoding *Ruminococcus gnavus*, PT-1 was drastically shifted to a *Bifidobacterium*-enriched, full-term-like community structure in media containing 2’FL. Our findings show that 2’FL, if supplemented into functionally matching microbiomes, could exert similar bifidogenic effects as HM in shaping gut microbiomes. This work led us to propose that glycosidase repertoires, as well as their cellular localization, should be investigated when assessing complex carbohydrate metabolism.

Our research uncovered strategies for enhancing the colonization of autochthonous *B. breve*, and potentially other *Bifidobacterium* spp., in preterm infants. Stable colonization of *Bifidobacterium* is crucial in early-life gut microbiome assembly and immune system development^71,72^. Compared to those of full-term, preterm infant gut microbiomes are depleted of persisting *Bifidobacterium* strains^37^. Besides clinical factors such as frequent antibiotic exposures and lack of sufficient breastfeeding^73,74^, we contend that microbiome compositions and functions, such as those dominated by Proteobacteria that cannot break down complex saccharides^37,43^, could also negatively affect the colonization of *Bifidobacterium*, especially those with a limited HMO metabolic capacity including *B. breve*. Thus, administering probiotics carrying complementary functions, such as extracellularly breaking down 2’FL into lactose and fucose, can be one approach to promote the growth and stable colonization of native *Bifidobacterium* in preterm infants.

Our work demonstrates the potential importance of personalized microbiome interventions. Specific probiotics and/or prebiotics may reshape gut microbiomes of distinct infants differently, thus influencing the subsequent gut microbiome assembly as well as infant development^75^. Widely prescribed probiotics show some positive outcomes, including reducing the chance of developing necrotizing enterocolitis (NEC) and sepsis in preterm infants^76,77^, yet their efficacies often vary^24,75^. For instance, *B. breve* is a commonly used probiotic^78^. However, partly due to its inability to metabolize many HMOs including fucosylated saccharides such as 2’FL, varied outcomes in infants have been reported^79–81^. Such variability also exists when using multi-strain probiotic cocktails. For instance, in a recent clinical trial, less than 40% of the preterm infant gut microbiomes responded positively to a five-strain probiotic treatment^82^. While other clinical factors such as diets and the immune system could affect the efficacy of microbiome interventions^75^, we experimentally showed that the resident gut microbiome represents a key factor in determining the treatment outcome. As shown with the preterm gut microbiome here, only when consortium-matching prebiotics and/or probiotics are administered, the expected health-promoting outcomes can be achieved.

In summary, by cultivating different infant gut microbiomes, we showed that community compositions influence HMO metabolism. Our work demonstrates the advantages of person-specific laboratory microbiomes that, when combined with metagenomics, may allow the identification of mechanisms shared across compositionally distinct microbiomes. The microbiome cultivation systems established here can be extended to experimentally elucidate individualized responses to other perturbations, such as antibiotics and other complex carbohydrates. The model microbiomes developed in this work can also be used when developing, testing, and using *in situ* microbiome editing tools to study mechanisms of inter-organism interactions in communities^83^. Over time, insights gained from these individualized *in vitro* microbiomes can lead to the development of precise, effective, and personalized microbiome treatments.

## Acknowledgments

We thank Michael J. Morowitz and Brian A. Firek for collecting infant stool samples characterized in prior work^37^; Rohan Sachdeva, Jordan Hoff, and Shufei Lei for their technical support; Netravathi Krishnappa and the rest of the IGI sequencing core for sequencing support; Steven A. Frese and Alexander Crits-Christoph for experimental suggestions; Matthew R. Olm for comments on the manuscript. For funding support, we acknowledge NIH award RAI092531A.

## Author contributions

Y.C.L., B.E.R., A.L.B., J.A.D., and J.F.B. designed the study; Y.C.L., K.D., B.E.R., R.R., and L.S. conducted experiments; Y.C.L., K.D., B.E.R., and R.R. performed DNA extractions and metagenomics library preparations for all samples; N.K. performed amplicon library preparation and sequencing; M.C.S. assisted with functional enrichment analyses; Y.C.L performed data analyses; Y.C.L and J.F.B. wrote the manuscript, and all authors contributed to the manuscript revisions.

## Resource availability

### Lead Contact

Further information and requests for resources should be directed to the Lead Contact, Jillian F. Banfield.

### Materials Availability

This study did not generate new unique reagents.

### Data and Code Availability

All data necessary for evaluating the conclusions in the manuscript are present in the manuscript and/or the supplementary files. The software used in this manuscript is publicly available. Reads of the stool inocula are available under BioProject PRJNA698986; SRA studies SRP304602; and SRA accessions SRS8184257 (FT-1), SRS8184183 (FT-2), and SRS8184427 (PT-1). Metagenome-assembled genomes will soon be available on NCBI and ggKbase (https://ggkbase.berkeley.edu/).

## Methods

### Human subjects

Refer to Lou et al.^37^ for detailed information on infant enrollment and stool collection.

### Bacterial strains

Whole genome sequences of *B. breve* and *R. gnavus* strains isolated from FT-1 will soon be available on NCBI and ggKbase (https://ggkbase.berkeley.edu/).

### Anaerobic growth media preparation

Aerobic Brain Heart Infusion (BHI) (BBL 211059), BHI without dextrose (Bioworld 306200141), and modified Gifu Anaerobic Broth (mGAM) (M2079-500G) were prepared using Milli-Q water as instructed by the suppliers. For preparing mucin-supplemented BHI media, we added 0.4% or 0.6% (wt/vol) mucin (type III mucin from porcine stomach; Sigma-Aldrich M1778) to BHI powder before mixing with Milli-Q water. Aerobic human milk (HM) (pooled from multiple donors; Lee Biosolutions, Inc) was prepared by centrifuging at 5,000 rcf for 5 min to remove most lipids.

Subsequently, all aerobic media solutions were sparged with N_2_ to remove oxygen and supplemented with 0.5 g/L L-cysteine hydrochloride and 1 mL/L of 0.1% resazurin sodium salt as a reductant and oxygen indicator, respectively, prior to autoclaving.

To prepare carbohydrate stock solutions (2’FL, L-fucose, and D-lactose), we dissolved 10g of the respective carbohydrate in 100 mL Milli-Q water and vacuum filtered (Corning™ 431475). Sugar solutions were loosely capped and stored in the anaerobic chamber for at least 48 hours before mixing with BHI-based media for cultivation.

### Infant gut microbiome cultivation

All cultivation work was performed in the same anaerobic chamber (Coy Labs; 70% N_2_, 25% CO_2_, 5% H_2_). The infant stool inoculum, stored in the −80°C freezer, was resuspended in PBS (phosphate-buffered saline) in a 1:2 wt/vol ratio. The resulting stool resuspension was homogenized by pipetting, and 15 μL of this mixture was added into 3 mL of a growth medium in a 24-deep-well block. The culture was allowed to recover for 48 hours at 37 °C with shaking, after which it underwent three more passages of 30 μL into 3 mL of fresh liquid medium, with each allowed to grow for 48 hours at 37°C. A cell-free negative control was included for each growth medium. After the first and the third (final) passaging, a 500-μL aliquot of the culture was taken for making glycerol stock, and a 1.5-mL aliquot was pelleted for DNA extraction and metagenomic sequencing. The remaining culture was used for pH measurements using a handheld pH meter (Apera model PH60S with the Swiss Spear pH Electrode (±0.01 pH accuracy)).

### DNA extraction, library preparation, and metagenomic sequencing for cultivated microbiomes

DNA extraction of the frozen cultivated microbiomes and/or fecal samples was performed using the Qiagen DNeasy PowerSoil DNA Isolation Kit. Library preparation was performed using the NEBNext Ultra II FS DNA Library Prep Kit for Illumina. Final sequencing-ready libraries were visualized using the Agilent 2100 Bioanalyzer before pooling, and the final pooled library was quantified through qPCR using the KAPA Library Quantification Kit for Illumina. Sequencing was performed using Illumina NextSeq 1000/2000 150 paired-end sequencing lanes with 10% PhiX spike-in controls. Post-sequencing bcl files were converted to demultiplexed fastq files per the original sample count with Illumina’s bcl2fastq v2.20 software.

### Mono- and co-culture experiments

All work was performed in the same anaerobic chamber (Coy Labs; 70% N2, 25% CO2, 5% H2) as the microbiome cultivation experiments described above.

*Bifidobacterium breve* and *Ruminococcus gnavus* from FT-1 were isolated using Bifidobacterium-Selective Agar (BSA; catalog # AS-6423). Specifically, the BHI+0.6% mucin and HM enrichments were streaked from frozen glycerol stocks onto four BSA plates (two per enrichment type) and incubated at 37°C for 48 hours. Forty colonies (10 per plate) were selected, streaked, and incubated at 37°C for 48 hours on BSA plates two additional times. The full-length 16S rRNA gene was amplified via colony PCR using primers 27F (5’-AGRGTTYGATYMTGGCTCAG-3’) and 1391R (5’-GACGGGCGGTGWGTRCA-3’) (Taq DNA Polymerase with Standard Taq Buffer kit (catalog #: M0273S)). The thermocycling conditions used were as follows: 1 cycle of 95°C for 3 min, 30 cycles of 95°C for 30 sec, 55°C for 30 sec, and 72°C for 45 sec, and 1 cycle of 72°C for 10 min. The amplicon was sequenced using Sanger sequencing, and colony taxonomies were identified using BLASTN against the 16S rRNA sequences from the NCBI BLAST Databases (downloaded in September 2022). Five colonies from each strain (*B. breve* and *R. gnavus)* were grown in MRS and BHI+0.6% mucin media, respectively, for 48 and 24 hours, respectively, before suspending in 50% glycerol and were stored at −80 °C. For all subsequent experiments, to ensure the consistency of the data, we used only one colony from each strain. The remaining four from each strain were stored as backups.

Prior to the mono- and co-culture experiments, *B. breve* and *R. gnavus* were inoculated from glycerol stocks and were grown anaerobically in 6 mL MRS and BHI+0.6% mucin media, respectively, for 48 and 24 hours, respectively. The cells were washed twice using anaerobic PBS by centrifuging at 6000 rpm for 10 min, the supernatants were discarded, and the cells were resuspended in 1.5 mL of PBS. OD_600_ was measured for the resuspended cultures. For the monoculture experiments, we inoculated an OD_600_ of 0.1 of the culture into the corresponding growth media (glucose-free BHI supplemented with 1% (wt/vol) of 2’FL, L-fucose, or D-lactose), and for the co-culture experiments, an equal OD_600_ of 0.05 of resuspended *B. breve* and *R. gnavus* cultures were inoculated into the same set of growth media as the monoculture experiments. All cultures were grown in triplicates for 144 hours, and OD_600_ was measured every 24 and/or 48 hours using Tecan microplate readers. Mono- and co-cultures were harvested 24, 72, and 144 hours after inoculation for community composition assessment. The mono- and co-culture growth rates were calculated as the slope of the natural log of the OD_600_ in the exponential phase over time.

### DNA extraction and amplicon sequencing for the mono- and co-culture experiments

DNA extraction of the mono- and co-cultures was performed via the DNeasy UltraClean 96 Microbial Kit. Relative abundance was estimated by amplicon sequencing. The V1–V2 regions of the 16S rRNA gene were amplified using a forward (5’-TGCTTAACACATGCAAGTCG-3’) and a reverse primer (5’-TCTCAGTCCCAATGTGGCCG-3’). The Phusion^®^ High-Fidelity PCR Master Mix (catalog #: M0531S) was used for PCR amplification. The thermocycling conditions used were as follows: 1 cycle of 98°C for 30 sec, 35 cycles of 98°C for 10 sec and 72°C for 10 sec, and 1 cycle of 72°C for 5 min. Amplicons were cleaned using 1.8X SPRIselect magnetic beads.

### *R. gnavus* microbiome supplementation assay

*R. gnavus* isolated from FT-1 and PT-1 enrichment passaged in BHI+0.4% mucin+0.3% 2’FL were inoculated from the glycerol stocks and were grown anaerobically in 6 mL BHI+0.6% mucin and BHI+0.4% mucin+0.3% 2’FL media, respectively, for 24 hours and 48 hours, respectively. The *R. gnavus* and the PT-1 enrichment glycerol outgrowths were then mixed in a 1:4 vol/vol ratio, and 30 uL of the resulting mixture was added into 3 mL of a growth medium in a 24-deep-well block. The remaining procedure followed the standard community cultivation assay (see the Method section “**Infant gut microbiome cultivation**”).

### Metagenomic assembly and gene prediction

Reads from all enrichments were trimmed using Sickle (www.github.com/najoshi/sickle), and those mapped to the human genome with Bowtie2^84^ under default settings were discarded. Subsequently, reads from each sample were assembled independently using IDBA-UD^85^ under default settings. Co-assemblies were also performed for each infant sample, in which reads from all enrichments of that infant sample were combined and assembled. Scaffolds that are <1 kb in length were discarded. The remaining scaffolds were annotated using Prodigal^86^ to predict open reading frames using default metagenomic settings.

### Metagenomic *de novo* binning

Pairwise cross-mapping was performed between all enrichments from each infant sample to generate differential abundance signals for binning. Each sample was binned independently using three automatic binning programs: metabat2^87^, concoct^88^, and maxbin2^89^. DasTool^90^ was then used to select the best bacterial bins from the combination of these three automatic binning programs. The resulting draft genome bins were dereplicated at 98% whole-genome average nucleotide identity (gANI) via dRep (v3.2.2)^91^, using minimum completeness of 75%, maximum contamination of 10%, the ANImf algorithm, 98% secondary clustering threshold, and 25% minimum coverage overlap. Genomes with gANI ≥95% were classified as the same species, and the genome with the highest score (as determined by dRep) was chosen as the representative genome from each species.

### Taxonomy assignment

The amino acid sequences of predicted genes in all assembled bins were searched against the UniProt100 database using the *usearch ublast* command with a maximum e-value of 0.0001. tRep (https://github.com/MrOlm/tRep/tree/master/bin) was used to convert identified taxonomic IDs into taxonomic levels. Briefly, for each taxonomic level (species, genus, phylum, etc.), a taxonomic label was assigned to a bin if >50% of proteins had the best hits to the same taxonomic label. GTDB-Tk was used to resolve taxonomic levels that could not be assigned by tRep^92^.

### Read-mapping-based species detection

Reads from each enrichment were mapped to the 1005 representative subspecies (generated from^37^). inStrain (v1.5.1) *profile*^93^ was run on all resulting mapping files using a minimum mapQ score of 0. Genomes with ≥0.5 breadth (meaning at least half of the nucleotides of the genome are covered by ≥1 read) with a minimum 0.1% relative abundance (number of reads mapped to the given genome divided by the total number of paired-end reads) in samples were considered present. If organisms were not detected in the initial inocula and were present in <40% of the replicates across all tested growth conditions, they were defined as contaminants and were removed from the final data tables.

### Pairwise genomic comparison of *B. breve* from the PS, S2, and S3

Genome-wide average nucleotide identity (gANI) for the three *B. breve* genomes reconstructed from the PS, S2, and S3 were calculated using dRep *compare* with the ANIm algorithm.

### Genome metabolic annotation

Kyoto Encyclopedia of Genes and Genomes (KEGG) orthology groups (KOs) were assigned to predicted ORFs for all fecal metagenomes using KofamKOALA^94^. Carbohydrate active enzymes (CAZymes) were assigned to all nucleotide sequences using run_dbcan.py (https://github.com/linnabrown/run_dbcan) against the dbCAN HMM (v11), DIAMOND (v2.0.9), and eCAMI databases with default settings. Final CAZyme domain annotations were the best hits based on the outputs of all three databases. Domains were also predicted using hmmsearch (v.3.3.2) (e-value cut-off 1 × 10^-6^) against the Pfam r35 database^95^. The domain architecture of each protein sequence was resolved using *cath-resolve-hits* with default settings^96^. The transporters were predicted using hmmsearch (same settings as the Pfam prediction and domain architecture was resolved using *cath-resolve-hits*) and BLASTP (v2.12.0+) (keeping the best hit, e-value cutoff 1e-20) against the Transporter Classification Database (TCDB) (downloaded in October 2022)^97^. SignalP (v.5.0b) was used to predict proteins’ putative cellular localization^98^.

### Metagenomics prediction on *B. breve’s* 2’FL utilization

Fucosidases GH29 and GH95, as well as their cellular locations, were searched on annotated *B. breve* genomes (see Method section “**Genome metabolic annotation**”). To search for potential 2’FL transporters, we first examined ten genes upstream and downstream of the identified *B. breve* GH95 fucosidase. No transporter hits were identified using a combination of the TCDB and KEGG databases. We next searched for homologs of two well-characterized *Bifidobacterium* fucosyllactose transporters using BLASTP^99,100^. All *de novo* assembled contigs from enrichments were used to account for potential mis-binning of the reconstructed *B. breve* genomes. The closest hit shares ~44% amino acid identity with the reference transporters, and its neighboring genes are not annotated as being involved in fucose- or lactose-related metabolism. We further mapped sequencing reads from enrichments against the reference fucosyllactose transporters and no hits were detected.

### Metagenomics prediction on D-lactose and L-fucose metabolism

To assess D-lactose and L-fucose metabolism, we first manually curated a list of relevant genes based on extensive literature searches^67,101–106^ (Table S5). For an organism predicted to metabolize D-lactose, it must encode lactose transporter(s) and β-galactosidase(s). For L-fucose metabolism, organisms must encode fucose transporter(s) and all genes involved in one of the L-fucose metabolic pathways (as listed in ^67,103^) to be identified as capable of L-fucose metabolism. Predicted pathways for each species were manually verified.

### Community diversity analysis

Modules from scikit-bio were used for the weighted (“skbio.diversity.beta.weighted_unifrac”) and unweighted (“skbio.diversity.beta.unweighted_unifrac”) UniFrac distances. The phylogenetic tree used in UniFrac distances was constructed by comparing all bacterial genomes grown in enrichments to each other using dRep *compare* with a mash sketch size of 10,000.

### Principal components analysis

PCA (Principal components analysis) was performed using *scikit-learn* and was conducted based on the relative abundance of bacterial genomes in enrichments as assessed using the weighted UniFrac distance.

### Quantification and Statistical Analysis

#### Two-group univariate comparisons

Statistical significance was calculated using Wilcoxon rank-sum test (implemented using the Scipy module “scipy.stats.ranksums”). Correlations were calculated using the Spearman correlation (implemented using the Scipy module “scipy.stats.spearmanr”). Multiple comparisons were false discovery rate (FDR) corrected with a threshold of q < 0.05.

#### Two-sided linear least-squares regression

For each infant stool inoculum, to calculate the reproducibility of the growth conditions, linear least-squares regression (implemented using the Scipy module “scipy.stats.linregress”) was applied to calculate the log_10_(relative abundance) of each species among pairs of replicates for all combinations of inoculum and medium.

