## Supplementary material for "Genomic analysis of cultivated infant microbiomes identifies *Bifidobacterium* 2’-fucosyllactose utilization can be facilitated by co-existing species": Figure S1

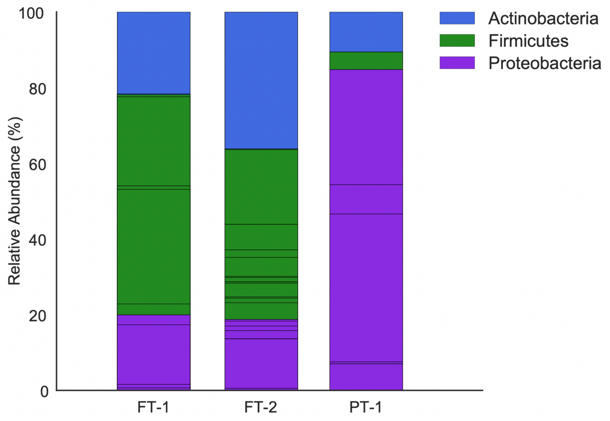


**Figure S1. Community compositions of the three infant stool inocula.** The x-axis represents the stool inocula. Bar height represents normalized species relative abundance, and bars are colored by phylum. Sections of the same color with horizontal black lines correspond to individual species of the same phylum.
