## Supplementary material for "Genomic analysis of cultivated infant microbiomes identifies *Bifidobacterium* 2’-fucosyllactose utilization can be facilitated by co-existing species": Figure S2

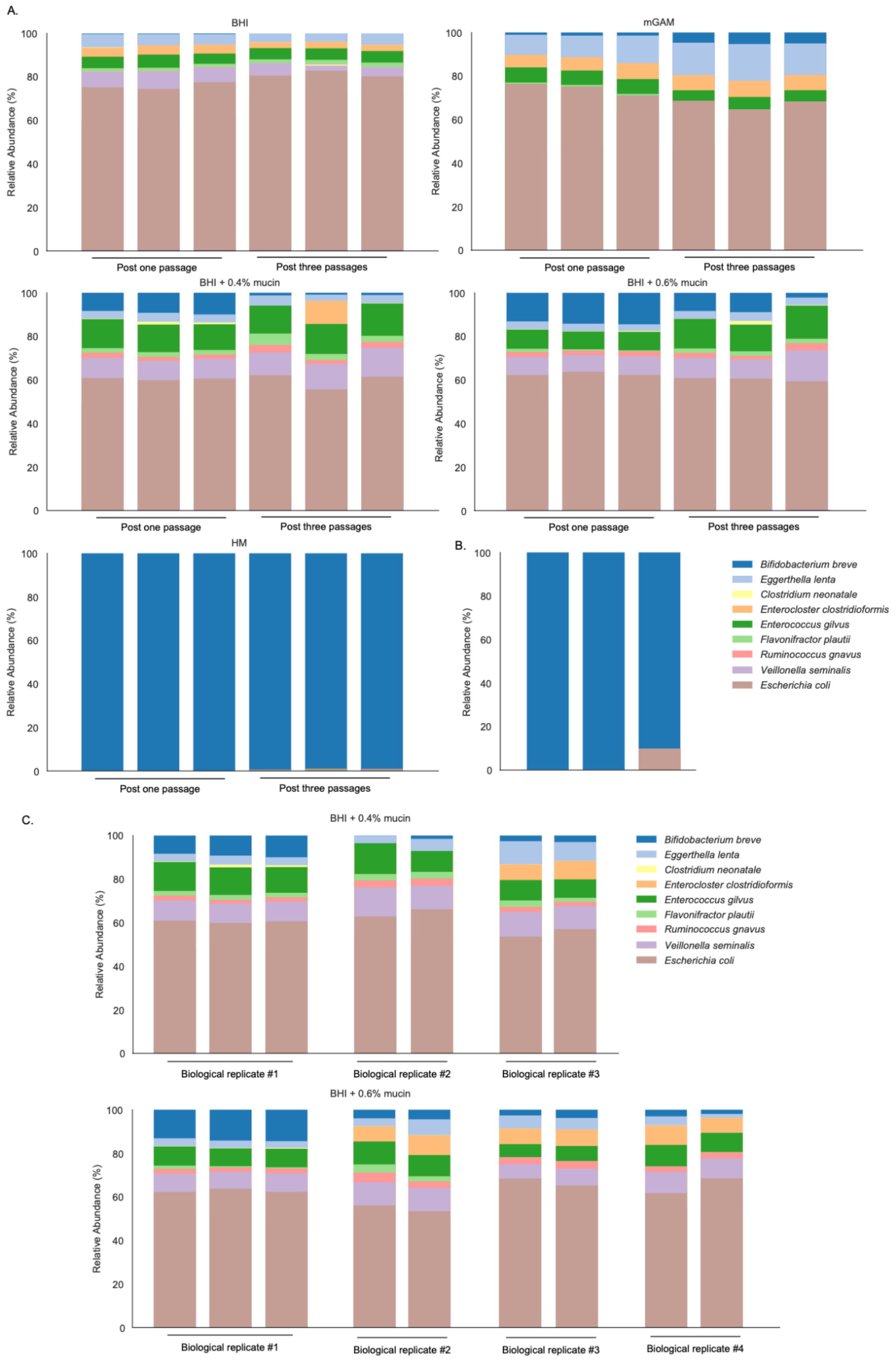


**Figure S2. Stability and Reproducibility of the FT-1 *in vitro* microbiomes.**

A) FT-1 in different growth media, and the compositions are compared between the first and the third passaging (x-axis, n=3). Bar height represents normalized species relative abundance, and bars are colored by species.
B) A HM-enrichment glycerol inoculum was grown in BHI + 0.6% mucin (n=3).
C) Reproducible FT-1 microbiome cultivation with BHI + 0.4% or 0.6% mucin. FT-1 inocula were inoculated in BHI + 0.4% or 0.6% mucin in three or four independent experiments (n=2 or 3).
