## Supplementary material for "Genomic analysis of cultivated infant microbiomes identifies *Bifidobacterium* 2’-fucosyllactose utilization can be facilitated by co-existing species": Figure S3

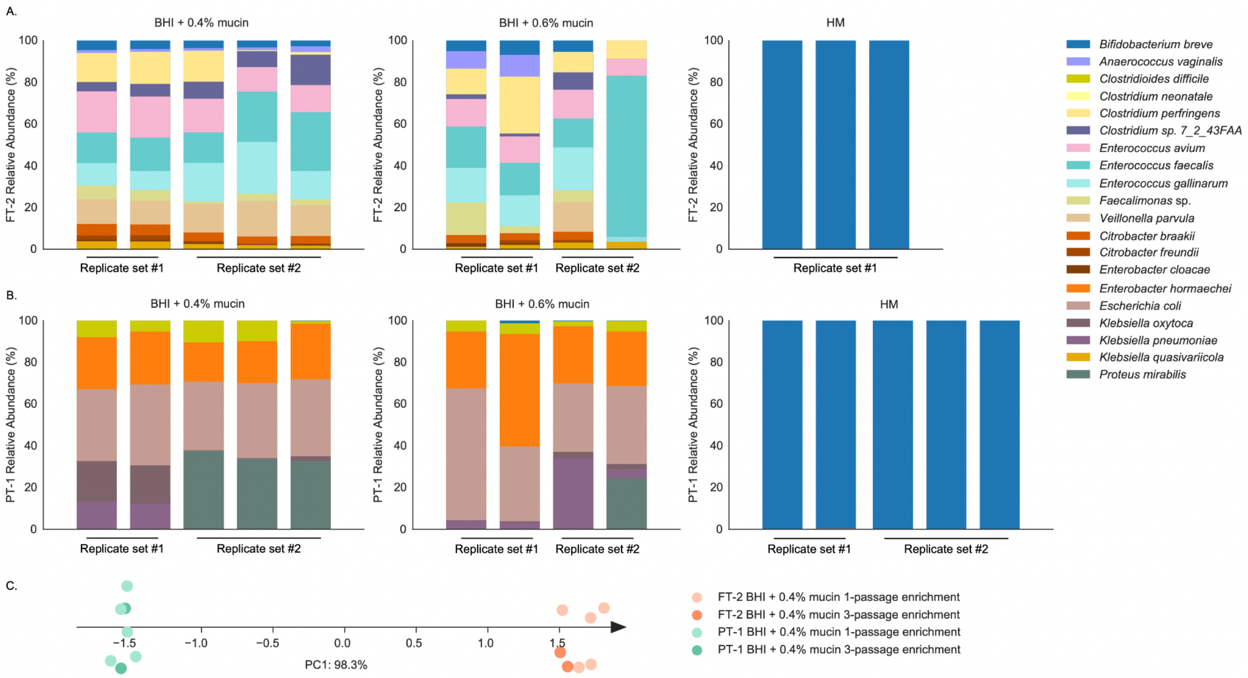


**Figure S3**. **FT-2 and PT-1 microbiome cultivation with BHI + 0.4% or 0.6% mucin**

A-B) S2 (A) and S3 (B) inocula were inoculated in BHI + 0.4% or 0.6% mucin. Bar height represents normalized species relative abundance, and bars are colored by species. The x-axis represents technical replicates.

C) PCA calculated using the weighted UniFrac Distances comparing FT-2 (green) and PT-1 (orange) communities grown in BHI + 0.4% mucin over three passages. Each circle represents a replicate.
