## Supplementary material for "Genomic analysis of cultivated infant microbiomes identifies *Bifidobacterium* 2’-fucosyllactose utilization can be facilitated by co-existing species": Figure S4

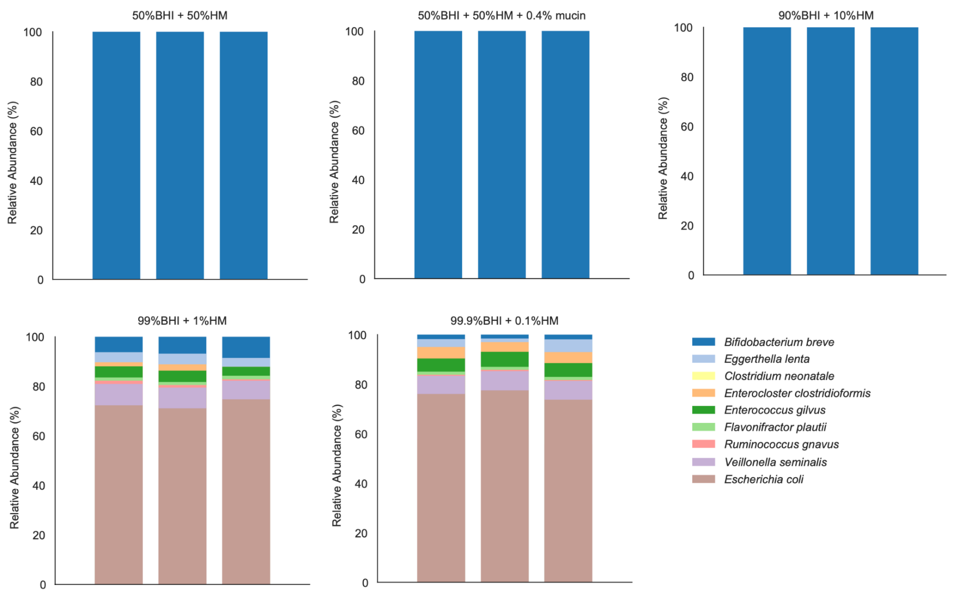


**Figure S4. FT-1 microbiome compositions in BHI-HM mixtures.** FT-1 inocula were inoculated in five BHI-HM mixtures. Bar height represents normalized species relative abundance, and bars are colored by species. The x-axis represents the technical replicates.
