## Supplementary material for "Genomic analysis of cultivated infant microbiomes identifies *Bifidobacterium* 2’-fucosyllactose utilization can be facilitated by co-existing species": Figure S5

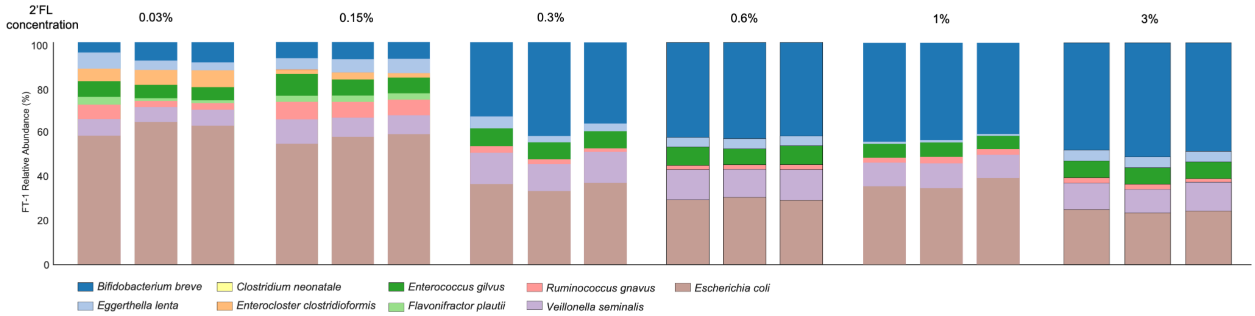


**Figure S5. The FT-1 microbiome grown in different concentrations of 2'FL**

The compositions of the FT-1 enrichments grown in BHI-mucin supplemented with varying concentrations of 2'FL (0.03%-3%) are shown in stacked bar plots. Bar height represents normalized species relative abundance, and bars are colored by species. The x-axis represents the growth media, and all replicates are shown.
