## Supplementary material for "Genomic analysis of cultivated infant microbiomes identifies *Bifidobacterium* 2’-fucosyllactose utilization can be facilitated by co-existing species": Figure S6

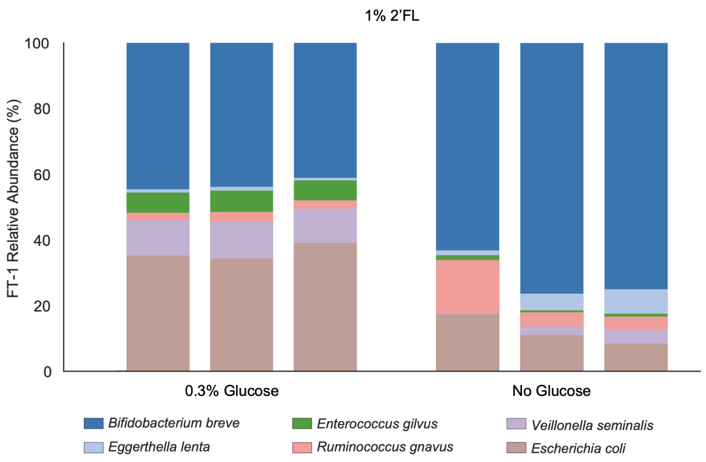


**Figure S6. Glucose concentrations influence the FT-1 microbiome composition**
The compositions of the FT-1 enrichments grown in BHI-mucin supplemented with 1% 2’FL with (left) and without (right) glucose. Bar height represents normalized species relative abundance, and bars are colored by species. The x-axis represents the growth media, and all replicates are shown. See also Table S4.
