## Supplementary material for "Genomic analysis of cultivated infant microbiomes identifies *Bifidobacterium* 2’-fucosyllactose utilization can be facilitated by co-existing species": Figure S7

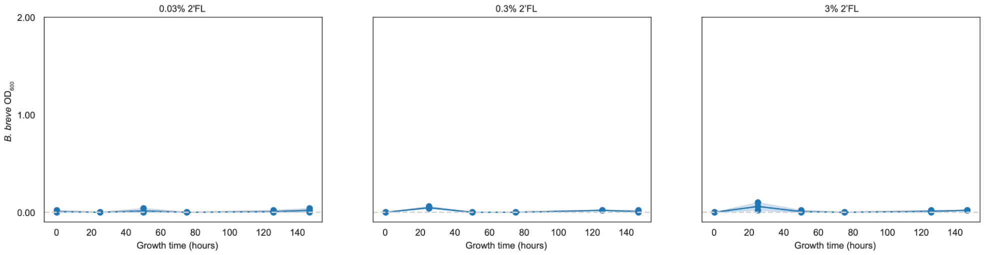


**Figure S7. *B. breve* monoculture growth in 2’FL.** *B. breve* (blue) grew in media in which 0.03% (left), 0.3% (middle), or 3% (right) 2’FL was added as the only carbohydrate source. The x-axis represents the growth time (in hours). The y-axis represents $OD_{600}$. Each circle represents a replicate.
